# Pathogen dependent phenotypic but no genetic correlation between sexual activity and 3 immunity in male *Drosophila melanogaster* subjected to differential sexual selection

**DOI:** 10.1101/144311

**Authors:** Zeeshan Ali Syed, Vanika Gupta, Manas Arun Samant, Aatashi Dhiman, Nagaraj Guru Prasad

**Affiliations:** Indian Institute of Science Education and Research Mohali, Sector-81, Knowledge city, SAS Nagar, Manauli PO. Mohali – 140306

**Keywords:** Sexual activity, Sexual selection, Immunity, trade-off, experimental evolution, drosophila

## Abstract

The theory of trade-off suggest that limited resources should lead to trade-off in resource intensive traits such as immunity related and sexually selected traits in males. Alternatively, sexual exaggerations can also act as an honest indicator of underlying immunocompetence, leading to positive correlations between these traits. Several studies have addressed this question using experimental evolution. However, they have rarely used ecologically relevant pathogens and fitness measurement (e.g., measuring post-infection survivorship) to find correlations between sexual selection and immunity. Here we attempt to address this caveat by evolving populations of *Drosophila melanogaster* under differential sexual selection. After more than hundred generations, we infected virgin and mated males from each population with three pathogenic bacteria: *Pseudomonas entomophila* (Pe), *Staphylococcus succinus* (Ss) and *Providentia rettgeri* (Pr). Fitness was measured as either post-infection survivorship (Pe and Ss) or bacterial clearance ability (Pr). Contrary to expectations, sexual selection had no evolutionary effect on male fitness against any of the pathogens. Moreover, mating had a beneficial effect against Pe and Pr, but no effect against Ss, suggesting pathogen specific phenotypic correlations between mating and immunity. Following these results, we discuss the significance of using ecologically relevant pathogens and quantifying host fitness while studying sexual selection-immunity correlations.

## Introduction

Two of the most important classes of traits that determine a male’s fitness are sexually selected traits, (i.e., traits which enable males to outcompete rivals in siring progeny) and, immunity related traits (i.e., traits which enable them to withstand pathogenic attack). Both sets of traits are resource intensive in their maintenance and deployment and, as life history theory suggests, are expected to trade-off with other life history related traits as a consequence (Stearns 1992). Traits such as longevity, stress resistance, fecundity etc. have been shown to trade-off with both immunity (Sheldon and Verhulst 1996; Moret *et al* 2000; Ye *et al* 2009; Ma *et al* 2012) and sexually selected traits (Johnston *et al* 2013; Nandy *et al* 2013). Such trade-offs are widespread, although not universal (e.g., Gupta *et al* 2016; Faria *et al*, 2015, Fricke and Arnqvist 2007), and are important in our understanding of why genetic variation exists in life history traits in the face of strong directional selection.

Following the idea of trade-offs, sexually selected and immunity related traits are also expected to trade-off with each other. Additionally, in males, such trade-offs can be apparent only with reproductive effort because several traits under sexual selection (such as courtship display and mating calls) manifest under the specific context of mating. Thus, populations undergoing differential levels of sexual selection might have differential effect of mating in their response to pathogenic infections. As an alternative possibility, to explain the results that deviated from the classical trade-off model, Hamilton and Zuk (1982) proposed that sexually selected traits in males might reflect their underlying immunocompetence, and therefore the two traits will be positively correlated. Studies addressing genetic correlation between mating and immunity in vertebrates have been the focus of much research following their pioneering work (Møller 1990; Balenger and Zuk 2014).

Due to the relatively simple immune system and small generation time of many invertebrate model organisms, it is possible to design tractable experimental evolutionary studies to test either hypotheses (Lawniczak et al. 2007). Phenotypic correlation between reproductive investment in males and several components of immunity has been studied in many invertebrate species. In wolf spiders, males presented with females increase their drumming rates at a cost of lytic activity (LA) (Ahtiainen *et al*. 2005). Negative correlations between encapsulation rate (EN), and, both call syllable number and spermatophore size were shown in bush crickets (Barbosa *et al*. 2016). In decorated crickets, artificial induction of spermatophore production traded off with phenoloxidase activity (PO) and LA (Kerr *et al*. 2010). Induction of immune system through lipopolysaccharide injection resulted in the reduction of their daily call rate (Jacot *et al.* 2005). In a more direct assay of immunological cost of mating, McKean and Nunney (2001) showed that increased sexual activity decreased the ability to clear non-pathogenic bacteria *E. coli* in male *Drosophila melanogaster*. Conversely, Gupta *et al*. (2013) found that mating increased ability to survive infection and clear the natural pathogen *Pseudomonas entomophila* in three unrelated populations of male *Drosophila melanogaster*. Similar results have also been found in bumblebees (Barribeau and Schmidt-Hempel 2016) and mealworm beetles (Valtonen *et al*. 2010).

Simmons *et al*. (2010) calculated quantitative genetic variation in immunity related and sexually selected traits in the Australian cricket *Teleogryllus oceanicus* using half-sib analysis and found negative genetic correlation between these two sets of traits. Mckean and Nunney (2008), using experimental evolution altered the intensity of sexual selection in laboratory populations of *Drosophila melanogaster* by skewing the sex ratio towards males. Higher sexual selection imposed on males resulted in lesser ability to clear the bacteria *E. coli*. In the yellow dung fly, *Scathophaga stercoraria*, removal of sexual selection through monogamy results in increased PO activity but that did not translate into increased antibacterial effect *in vitro* (Hosken 2001). In *Tribolium castaneum* similar removal of sexual selection did not result in difference in either PO or their ability to survive the infection by the pathogenic microsporidian *Paranosema whitei* (Hangartner *et al* 2015). Thus, the evolutionary relationship between sexually selected traits and immunity, at least in invertebrates is unclear.

A “recurring theme” in a lot of the above mentioned studies, as observed by Lawniczak et al. (2007), is the lack of a fitness oriented experimental framework. Differences in molecular readouts (such as gene expression, PO and LA) do not always translate into fitness differences (e.g., Hangartner *et al* 2015). This leads to a dissonance between potential immunity (gene expression, PO, LA etc.) and realized immunity (actual ability to survive pathogenic infection) (Fedorka *et al* 2007). One way of incorporating fitness is to evolve host populations under different levels of sexual selection, and then measure their fitness (e.g., survivorship) against pathogenic infection. That said, even the supposedly simple immune system of invertebrates is in fact not that simple, with several studies showing pathogen specificity (Sadd and Schmid-Hempel 2009), immune memory (Kurtz 2005), and transgenerational immune priming (Sadd *et al* 2005). The pathogen(s) that a host is being exposed to constitute an important part of the host’s ecological context and can play a non-trivial role in determining the outcome of the interaction between reproductive investment and realized immunity. If the same host responds through different immune mechanisms to different pathogens (i.e., specificity), mating may have differential effect on host ability to combat different infections. For example, Gupta *et al*. (2013) showed that males from the same populations of *Drosophila melanogaster* that showed increased resistance against *Pseudomonas entomophila* upon mating, did not show any effect of mating when challenged with *Staphylococcus succinus*. This argument can be extended to the evolutionary effect of sexual selection in males on their immune response as well. Therefore, in order to assess these relationships, it is important to measure host fitness against different ecologically relevant pathogens. However, such studies are conspicuous by their absence.

In this study we try to address this issue by evolving replicate populations of *Drosophila melanogaster* under increased and decreased levels of sexual selection for more than a hundred generations. Alteration of sexual selection was achieved by maintaining the populations under male biased (M) or female biased (F) operational sex ratio regimes. Previous studies have shown that males in these populations have diverged in terms of their reproductive traits, such as courtship and locomotor activity, and, sperm competitive ability (Nandy et al 2013 a,b). We subjected the males from both regimes to infection by three ecologically relevant bacteria *–Pseudomonas entomophila* (Pe), *Staphylococcus succinus* (Ss), and *Providencia rettgeri* (Pr) in three different assays. To address the effect of mating, in each of the assays, we had two groups of males from each selection regime – virgin and sexually active. We used survivorship post infection as a measure of fitness in two of the assays (Pe and Ss), and ability to clear bacteria in the third (Pr). We predicted that a common correlation between sexual selection and immunity (physiological and/or genetic) would produce similar differences in fitness between M and F males across all three assays, whereas pathogen dependency would produce different results for different pathogens.

## Materials and Methods

### Ancestral populations

The two ancestral populations used in this study are called LH and LH_st_, both of which are large laboratory adapted populations of *Drosophila melanogaster*. The LH population was established with 400 gravid wild caught females and is maintained at an adult census size >5000 (Chippindale and Rice 2001). LHst was derived by introgressing a benign autosomal ‘scarlet eye’ marker to the LH genetic background and is maintained at an adult census size >2500. The LH and LH_st_ populations are genetically equivalent except for one locus which has no discernible effect on their fitness (Bodhi thesis or some other ref?). The LH_st_ population is periodically back crossed with LH population to prevent divergence between the two populations (Prasad *et al*. 2007). Both populations are maintained at standard laboratory conditions (temperature = 25±1°c, relative humidity ≈ 60%, 12:12 dark: light cycle) on corn-meal molasses food. Detailed population maintenance is described in Nandy *et al*. (2012). Briefly, in a given generation, 2-3 day-old adult flies from rearing vials (95 mm height × 25 mm diameter) are mixed and redistributed into fresh food vials with 16 males and 16 females in each and containing a limiting quantity of dried yeast granules. The flies are kept there for two days after which they are allowed to oviposit for 18 hrs in fresh vials with food. These vials are controlled for egg density (~150 eggs /vial) and incubated to start the next generation.

### Selection regimes

The selection regimes are derived from LH_st_. Initially three populations, C_1-3_, were derived and maintained for 5 generations. The maintenance of the C populations differed from that of LH_st_ in that adult males and females were collected as virgins and held in same-sex vials with 8 individuals/vial and combined in 1:1 sex ratio (16 males and 16 females) once they were 2-3 days old with measured amount of live yeast instead of granules. Thereafter the maintenance protocol is the same as that of LH_st_. After 5 generations, two more selection regimes, F_1-3_ and M_1-3_, were derived from each of the C populations where operational sex ratios where biased towards males and females respectively. In these populations, 2-3 day-old virgin adults were combined in their respective sex ratios (i.e., 1Male:3Females for F populations and, 3Males:1Female for M populations). Note that the populations sharing the same subscript share a common ancestry and are handled simultaneously, independent of those having a different subscript. Thus each subscript constitutes a “statistical block”. Details of maintenance and selection history are described in Nandy et al (2013).

### Standardization

Nongenetic parental effects (Rose 1984) can lead to misinterpretation of multi-generation selection experiment results. To equalize such effects across selection regimes, all selected populations were passed through one generation of standardization where selection was removed, i.e., they were maintained in ancestral conditions (Syed et al 2016). Adult progeny produced by this generation were used for the experiment.

### Bacterial culture

We used three pathogens for this study: gram negative bacteria *Providencia rettgeri* (Short and Lazzaro 2013), gram negative bacteria *Pseudomonas entomophila* L48 (Vodovar *et al*. 2005), and gram positive bacteria *Staphylococcus succinus subsp. Succinus*, strain PK-1 (Ss) (Singh *et al*. 2016). All three bacteria are natural isolates obtained from wild caught *Drosophila*. For making the bacterial suspension for infections, bacterial culture was grown at 27°C (Pe) and 37°C (Ss and Pr) till OD☐1 = ☐1 1.0☐±☐0.1 from a glycerol stock maintained at -80°C. Following this, cells were pellet down and suspended in equal volume of 10 mM MgSO4 before infection. For Pr, the suspension was concentrated to OD 2.0☐±☐0.1 before infection.

### Infection protocol

Flies were put under light CO_2_ anaesthesia and infected by pricking with a needle (*Minutein pin* 0.1 mm, Fine Science Tools, CA) dipped in bacterial suspension (bacteria suspended in 10 mM MgSO4) in the thorax (Gupta et al 2013). To control for injury, a separate set of flies were pricked with a tungsten needle dipped in sterile 10 mM MgSO_4_ (sham).

### Experimental Treatments

Experimental males were collected within 6 hours of eclosion from pupae, which ensured their virginity, since in these populations it takes the flies ~8 hours to attain sexual maturity. These males were kept in vials provided with corn-meal molasses food at a density of 8 males per vial. On 12^th^ day post egg collection (i.e., 2-3 day-old adult) flies from each selection regime were randomly assigned to two groups: ‘Virgin’ and ‘Mated’.

In the Virgin treatment, virgin males were transferred to vials containing fresh food as they were. In the Mated treatment, males from each vial were combined with virgin LHst females (8 / vial). For infection and sham, 10 (n = 80) and 5 (n= 40) vials were set up per treatment per selection regime per block respectively for each of the pathogens. All pricking was done on 14^th^ day post egg collection and were transferred to vials containing fresh food following infection. Males in the ‘mated’ treatment were separated from females while anaesthetized for pricking and were maintained in single sex vials.

### Measure of infection response

For Pe and Ss response to pathogenic infection was measured in terms of survivorship post infection by observing vials for mortality every three hours post infection for ~100 hours post infection. For Pr, since mortality was low (<5%) and did not differ from the sham control, response was measured as the ability of the host to clear bacteria using the method described in Gupta et al (2013). Briefly, 20 hours post infection, 6 flies from each vial was sampled randomly and divided into groups of three. They were then crushed using a mortar inside micro-centrifuge tubes containing 100 μL MgSO_4_ and plated on LB-Agar plates using an automated spiral plater (WASP spiral plater, Don Whitley Scientific, UK). Three replicate plates were plated from each group of three flies. After growing the bacteria in their respective optimum temperatures, CFUs were counted using a plate reader (Acolyte colony counter, Don Whitley Scientific, UK). Average CFUs per fly obtained from each group was used as unit of analysis.

### Statistical Analyses

All analyses were performed in R (R core team 2015). Survivorship (for Pe and Ss) was analysed using Cox’s Proportional hazards model. Death was recorded for each fly and flies not dead till the last time were treated as censored data. For each of the pathogens, data were modelled either using block as a random factor or excluding Block using R package “Coxme” (Therneau 2012) using the following two expressions:

> Model 1: ~Selection_Regime * Mating_Status + (1 | Block/Selection_Regime)
>
> Model 2: ~Selection_Regime * Mating_Status

Since analysis of deviance revealed no effect of block (analysis of deviance test: 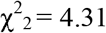, p = 0.12 for Pe; 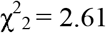, p = 0.27 for Ss), data from all three blocks were pooled and the cumulative data were then tested for difference in survivorship. We compared the Cox partial likelihood (log-likelihood) estimates across treatments and selection regimes.

Test for the effect of mating status and selection on number of CFUs (in case of Pr), colony count data was natural log transformed. Normality was verified using Shapiro – Wilk test. The data were then subjected to ANOVA (with Satterthwaite approximation for degrees of freedom) where mating status and selection regimes were used as fixed factors and block as a random factor. This was performed using package “lme4” (Bates et al. 2011).

Post-hoc pairwise comparisons using Tukey’s HSD method was performed with the package “lsmeans” (Lenth 2016). All data are archived in Dryad data repository.

### Results

For survival analysis, we compared the Cox partial likelihood (log-likelihood) estimates. Mating had a significant effect on survival against Pe (Table1a). Pairwise comparisons showed that mated males survived better than virgins in both M and F regimes (p<0.001, Figure 1). However, there was no effect of selection or selection × mating status interaction. This indicates that sexual selection had no effect on survival on either virgin or mated males. Similarly, against Pr, we found a significant effect of mating, but no selection × mating interaction effect (Table 1c). Here again, post-hoc analysis (Tukey adjusted p) showed that mated males were able to clear more bacteria compared to virgins in both M (p= 0.045) and F (p=0.015) regimes (Figure 2). There was no effect of either mating, selection regime or selection × mating interaction on survivorship against infection by Ss (Figure 3, Table 1b).

**Table 1:** Analysis of cox proportional hazards for survivorship post-infection for *A. Pseudomonas entomophila* and (B) *Staphylococcus succinius*. (C) Analysis of bacterial colony count data (natural log transformed) against *Providencia rettgeri*.

| A. Survivorship against *Pseudomonas entomophila* |  |  |  |  |
| --- | --- | --- | --- | --- |
|  | loglik | Chisq | df | Pr(>\|Chi\|) |
| Selection | -5035.2 | 0.9701 | 1 | 0.325 |
| Mating | -5011.8 | 46.8518 | 1 | 7.656e-12* |
| Selection × Mating | -5011.6 | 0.2909 | 1 | 0.59 |
| B. Survivorship against *Staphylococcus succinus* |  |  |  |  |
| Selection | -4377.6 | 3.4314 | 1 | 0.064 |
| Mating | -4377.6 | 0.0556 | 1 | 0.813 |
| Selection × Mating | -4377.5 | 0.1624 | 1 | 0.687 |
| C. Bacterial clearance ability against *Providencia rettgeri* |  |  |  |  |

|  | Sum Sq | Mean Sq | NumDF | DenDF | F.value | Pr(>F) |
| --- | --- | --- | --- | --- | --- | --- |
| Selection | 0.447 | 0.447 | 1 | 4.7040 | 0.0986 | 0.766948 |
| Mating | 60.619 | 60.619 | 1 | 15.3587 | 13.3857 | 0.002249* |
| Selection × Mating | 0.126 | 0.126 | 1 | 7.0301 | 0.0279 | 0.872035 |

**Figure 1:**
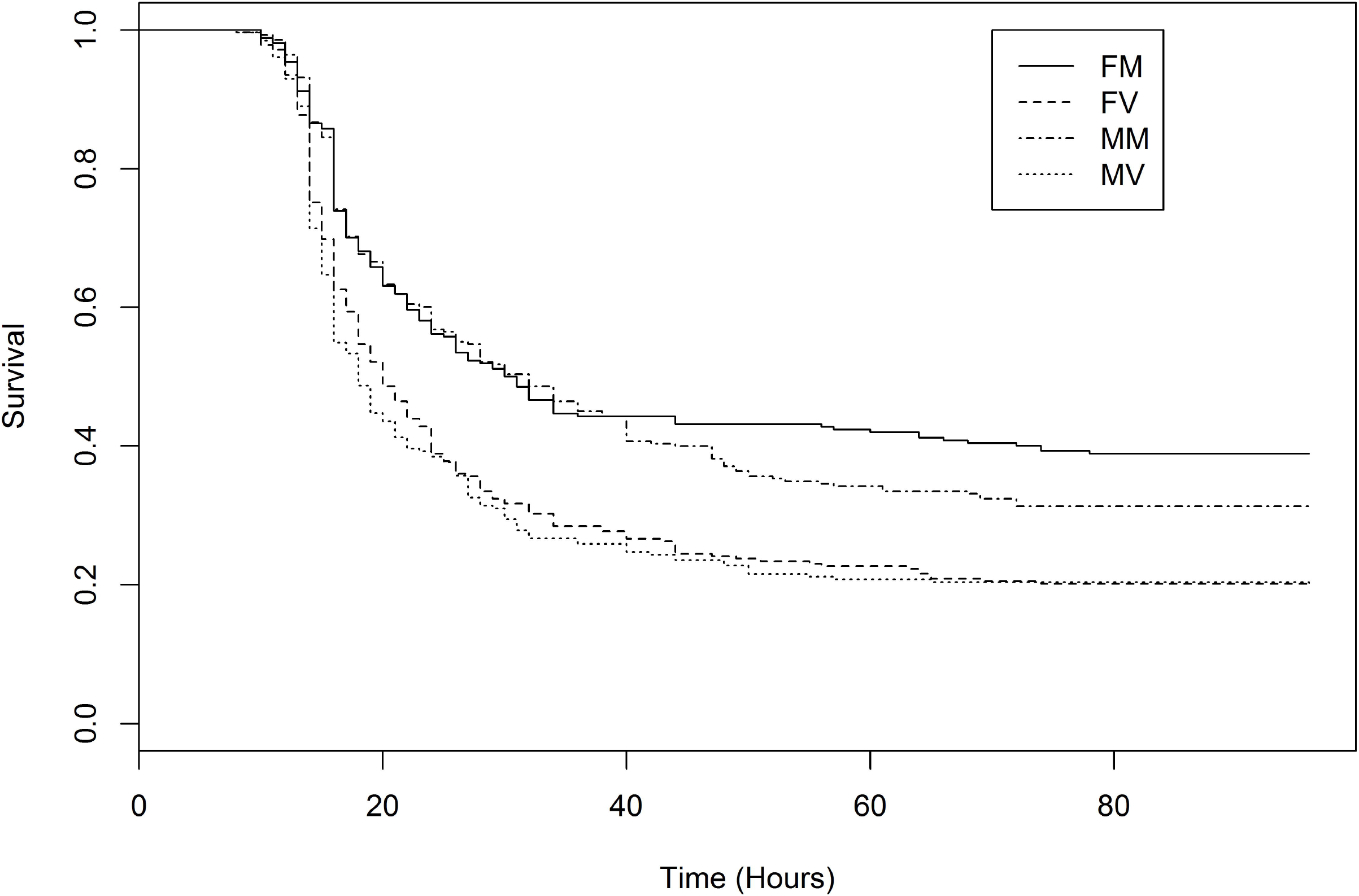
Results of Cox proportional hazards analysis. The curves show survival as a function of time. The solid line and the dot-dash line represent “mated” treatments of F (FM) and M (MM) regimes respectively. They have significantly higher survival rate compared to the “virgin” treatments of F (FV) and M (MV) regimes denoted by dashes and dots respectively.

**Figure 2:**
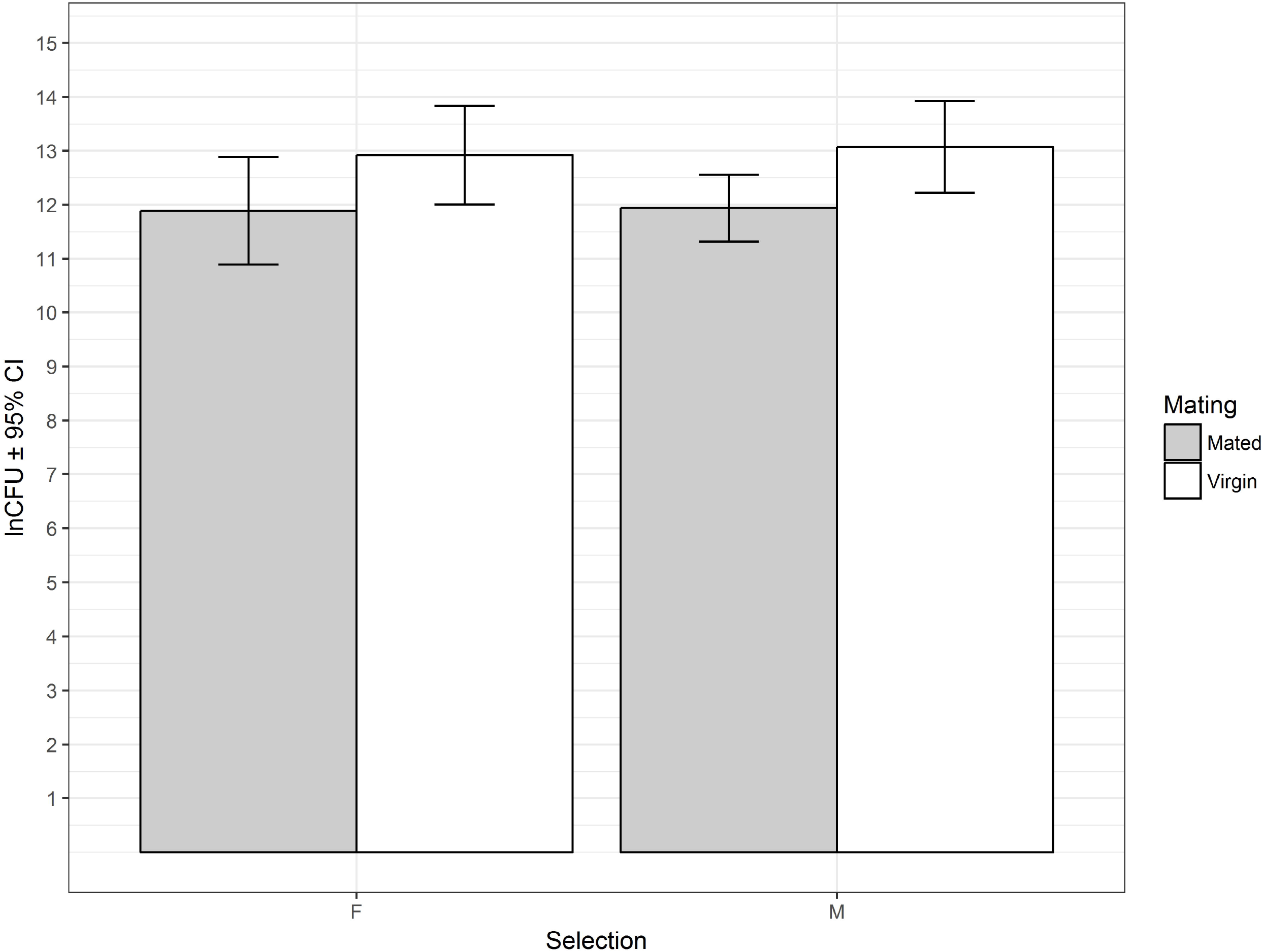
Results of natural log transformed CFU data for mated (Shaded bar) and virgin (open bar) treatments of M and F regimes which are represented in the x-axis. The error bars represent 95% confidence intervals. In both selection regimes mated males had significantly lower colony count than virgins.

**Figure 3:**
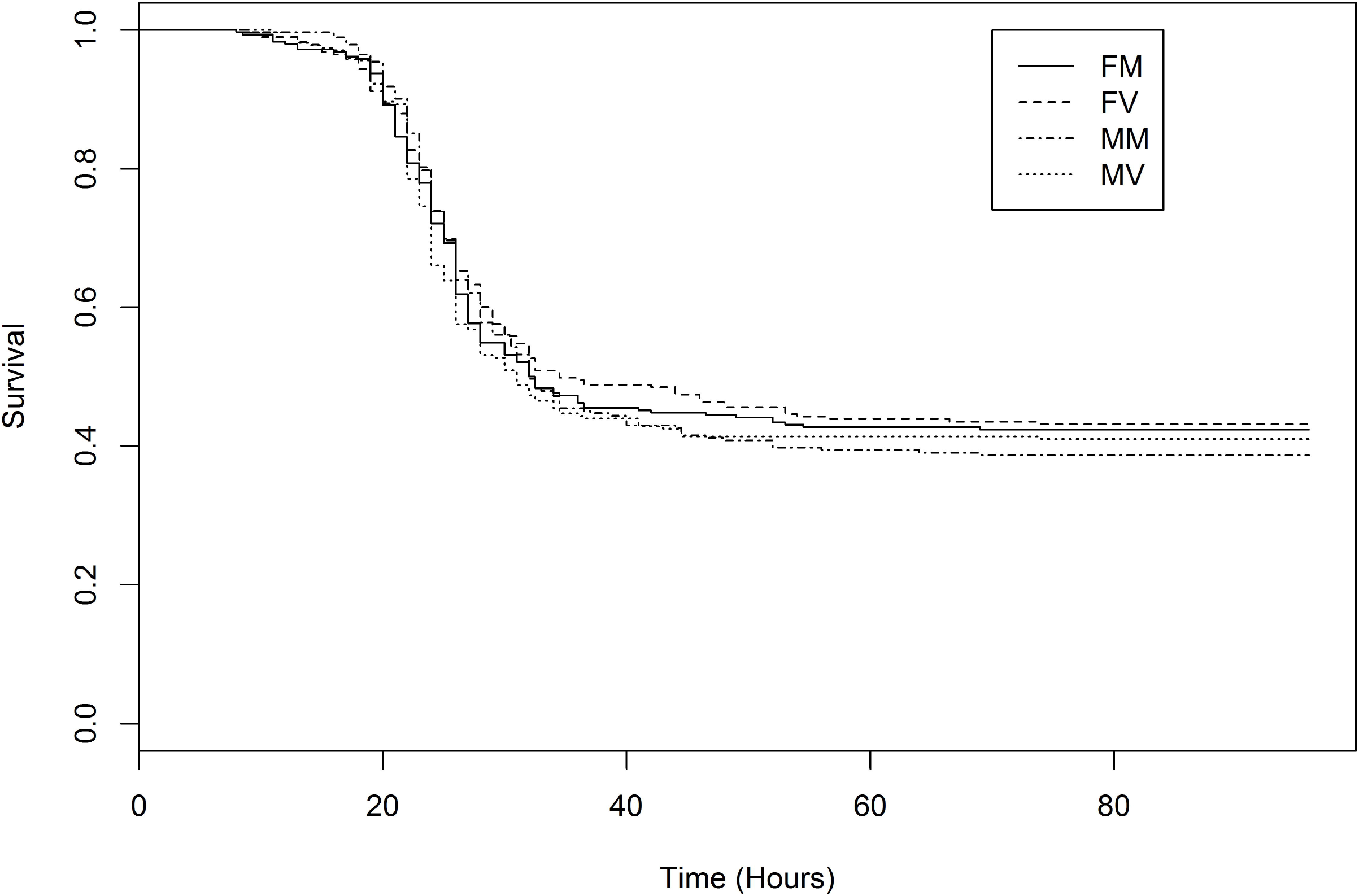
Results of Cox proportional hazards analysis. The curves show survival as a function of time. The solid line, the dot-dash, the dashes and the dots line represent “mated” treatments: F-Mated (FM), M – mated (MM), F-virgin (FV) and - virgin (MV) respectively. There was no differences between any of the treatments.

#### Discussion

Here we show, using *Drosophila melanogaster* and three different ecologically relevant pathogens – *Pseudomonas entomophila* (Pe), *Providencia rettgeri* (Pr) and *Staphylococcus succinus* (Ss), that mating affects the host’s ability to survive or clear bacterial infection in a pathogen dependent manner. Further, we used experimental evolution in the same system to show that even after going through more than a hundred generations of differential sexual selection, populations of *Drosophila melanogaster* did not differ in either their ability to fight pathogenic infection or the effect of mating on their ability to fight pathogenic infection.

#### No effect of sexual selection on immune response

Males from M and F regimes were not different from each other in terms of either post infection survivorship (against Pe and Ss) or bacterial clearance ability (against Pr), given that they were from the same mating treatment (mated or virgin). Our results differ from that of McKean and Nunney (2008), who measured host’s ability to clear *E.coli* as a proxy for immune response, and found a trade-off with sexual selection. This shows that relations between multi-locus traits such as immunity related traits and traits under sexual selection can be complex and may not follow a simple “Y-model” of trade-off (Stearns 1992). Several other studies have measured one or a few components of immunity, such as phenoloxidase activity and found them to be negatively correlated with the intensity of sexual selection (Hosken 2001, Hangartner *et al* 2015). However, such studies that measure one (or a few) component of immunity to assay the effect of sexual selection on immunity can have certain drawbacks. Different components of the immune system can have their own internal correlations. For example, negative genetic correlation between resistance and tolerance has been reported in a mice-Plasmodium *chabaudi* system (Råberg *et al*. 2007). Within-immune system trade-offs have also been found in female white-footed Mice, *Peromyscus leucopus* (Martin *et al*. 2007). Therefore measuring just one or a few of them can lead to incomplete and perhaps misleading conclusions about the genetic correlations between immunity and sexual selection. Furthermore, some of these components might have no fitness consequence. Leclerc *et al*. (2006) found that in *Drosophila melanogaster*, mutants that failed in producing active phenoloxidase had equal survivorship compared to wild type flies against pathogenic infection by different species of fungi and, both gram positive and negative bacteria. This implies that in flies, phenoloxidase activity is probably unnecessary for survival against a wide variety of microbes (Leclerc *et al*. 2006). Similarly, Hosken (2001) found that removal of sexual selection resulted in increased PO activity in males which, however did not result in increased antimicrobial activity. Thus, while measuring components of immunity is important for understanding the functional basis, it’s the fitness consequence that ultimately drives the evolution of a trait, and it is therefore important to measure immunity in that context. In the present case, we have used three different natural isolates of *D. melanogaster* and showed that neither survival nor bacterial clearance ability changes in response to differential levels of sexual selection, providing persuasive evidence that in this system response to sexual selection has not been traded-off with investment in immune response.

#### Phenotypic effect of reproduction on immunity depends on pathogen

We found that mated males from both M and F regimes had better survivorship and bacterial clearance abilities against Pe and Pr respectively. We have previously shown that in the population ancestral to the selection lines used here, mating had a beneficial effect on resistance against Pe (Gupta *et al*. 2013). Our results corroborate with those of Valtonen *et al*. (2010) and, Barribeau and Schmidt-Hempel (2016) who also found that mating can be beneficial against infections. However they are in contrast with McKean and Nunney (2001) in that, unlike this study, we found no evidence of trade-off between mating and immunity in terms of either survival, or bacterial clearance. Phenotypic relationships between multi-component traits such as immune response (with mutually non-exclusive components such as resistance, tolerance, memory etc.) and reproduction (with components such as acquisition of mates, production of sperm and accessory gland proteins etc.) is complex – even invertebrates like fruit flies show great variety and pathogen specificity in their response to infections. Thus, measuring such relationships is expected to depend upon the pathogens. The fact that we find no difference in survivorship between mated and virgin males against a third pathogen, Ss (flies used randomly from the same experimental pool, see methods.), further highlights this issue.

#### Evolutionary response does not mirror phenotypic correlation

McKean and Nunney (2008) showed that increased sexual selection resulted in evolved populations of *Drosophila melanogaster* where males had exaggerated sexually selected traits, but had reduced ability to clear the non-pathogenic bacteria *E. coli*. This mirrored the phenotypic trade-off they found between mating and immunity (McKean and Nunney, 2001, 2008). Our results differ from McKean and Nunney (2008) in that we did not find any phenotypic trade-off. Quite contrarily, for Pe and Pr we found that mated males had higher survivorship and bacterial clearance ability respectively. Thus, this did not mirror the genetic correlations, where males from both M and F regimes had equal ability to survive infection or clear bacteria within a given mating treatment. The most likely explanation is that, phenotypic correlations do not necessarily mirror genetic correlations (Chippindale *et al*. 1997). Expression of immunological reaction to infection and sexually selected traits (such as aggressiveness towards rival males, courtship display, production of sperm and accessory gland proteins) depend on the condition of the condition of the organism, which can be affected by factors such as age, developmental conditions, resource availability etc (Stearns et al., 1991). Therefore these factors might impact the correlations between these traits through Genotype × Environment interactions. For example, in *Drosophila melanogaster*, Genetic correlations between immunity and other life history related traits have been found to be dependent upon the host dietary condition (McKean *et al*. 2008) and temperature at which they were maintained (Lazzaro et al 2008). Therefore, it is possible the phenotypic and genetic relations between sexual selection and immune response might manifest in conditions that differ from their maintenance regime (such as resource limitation).

#### Summary

Using three different pathogens of *Drosophila melanogaster*, we found no evolutionary effect of the intensity of sexual selection on the immunocompetence of males. This is in contrast with several previous studies (Zuk and Stoehr 2002; Schmid-Hempel 2003, McKean and Nunney 2008). We also show that mating can have beneficial or no effect on males depending upon pathogen. This adds to a growing body of studies that have used natural pathogens to show the beneficial effects of mating on hosts (Valtonen *et al*., 2010, Gupta *et al*. 2013, Barribeau and Schmidt-Hempel, 2016). Taken together, our study indicates that the complex relationship between reproductive investment and immune response might not manifest in the form of simple trade-offs. We also highlight the importance of ecologically relevant pathogens and incorporating host fitness, such as survival ability against infection in measuring such relationships.

## Author contributions

SZA, VG, and NGP thought of the study. ZSA, VG, AD and MAS conducted the experiments and collected data, ZSA and VG analysed the data. All authors contributed in writing and reviewing the manuscript.

## Acknowledgments

The authors would like to thank Ms Radhika Ravikumar and Ms Saudamini Venkatesan for helpful comments on the manuscript; Mr. Nagendra Kumar for assistance with equipment and members of the Evolutionary Biology Lab, IISER Mohali for helping out with experiments and population maintenance. ZSA, MAS and VG thank Council for Scientific and Industrial Research, Govt of India for Junior and Senior Research Fellowships. AD thanks Department of Science and Technology, Govt. of India for INSPIRE fellowship. All authors thank IISER Mohali for lab and logistic support.

## Notes

**Conflict of Interest Statement:** The authors have no conflict of interest.

